# Common data models to streamline metabolomics processing and annotation, and implementation in a Python pipeline

**DOI:** 10.1101/2024.02.13.580048

**Authors:** Joshua M. Mitchell, Yuanye Chi, Maheshwor Thapa, Zhiqiang Pang, Jianguo Xia, Shuzhao Li

## Abstract

To standardize metabolomics data analysis and facilitate future computational developments, it is essential is have a set of well-defined templates for common data structures. Here we describe a collection of data structures involved in metabolomics data processing and illustrate how they are utilized in a full-featured Python-centric pipeline. We demonstrate the performance of the pipeline, and the details in annotation and quality control using large-scale LC-MS metabolomics and lipidomics data and LC-MS/MS data. Multiple previously published datasets are also reanalyzed to showcase its utility in biological data analysis. This pipeline allows users to streamline data processing, quality control, annotation, and standardization in an efficient and transparent manner. This work fills a major gap in the Python ecosystem for computational metabolomics.

**Author Summary:** All life processes involve the consumption, creation, and interconversion of metabolites. Metabolomics is the comprehensive study of these small molecules, often using mass spectrometry, to provide critical information of health and disease. Automated processing of such metabolomics data is desired, especially for the bioinformatics community with familiar tools and infrastructures. Despite of Python’s popularity in bioinformatics and machine learning, the Python ecosystem in computational metabolomics still misses a complete data pipeline. We have developed an end-to-end computational metabolomics data processing pipeline, based on the raw data preprocessor Asari [1]. Our pipeline takes experimental data in .mzML or .raw format and outputs annotated feature tables for subsequent biological interpretation. We demonstrate the application of this pipeline to multiple metabolomics and lipidomics datasets. Accompanying the pipeline, we have designed a set of reusable data structures, released as the MetDataModel package, which shall promote more consistent terminology and software interoperability in this area.

## Introduction

Metabolomics aims to comprehensively detect, identify, and quantify the diverse small molecules, i.e., metabolites, present in biological systems. This provides key information on biochemical phenotypes, often reflecting the function of genes and genomes. With the progress of technologies, metabolomics is becoming a regular component of many biomedical projects [2] [3] [4] [5]. Thousands of metabolomics datasets are now available in major data repositories [6] [7] [8] and the annual citation of “metabolomics” in PubMed now exceeds ten thousand. Due to this increasing popularity, solutions for processing such data need to be better incorporated into the regular bioinformatics workflows [9] [10] [11]. This integration will require an ecosystem in both the R and Python programming languages, the two dominant languages for bioinformatics, each with unique strengths and a large user community.

The foundational tool of a software ecosystem in computational metabolomics is the preprocessing tool that, among other functions, converts raw data into feature tables representing signals of interest likely to represent metabolites. XCMS [12] has served this role for the R programming language, and various tools for further data processing, including annotation, quality assurance and quality control (QA/QC), have been built utilizing its outputs [13] [14] [15] [16]. Many optimization tools and pipelines have been built around XCMS [17] [18] [19] [20] [21]. Despite the popularity of Python in machine learning and bioinformatics in general, a robust ecosystem for metabolomics in Python remains lacking, primarily due to the lack of a preprocessing tool for metabolomics raw data. While a handful of Python tools have been developed over the past decade [22] [23], they are either dated or not production-ready. With the recent release of Asari [1], a preprocessing tool implemented in Python, Python has become a viable option for data processing in computational metabolomics.

As computational metabolomics evolves, the community continues working to define operational terminology and best practices. These efforts have resulted in various workgroups and multiple publications [24] [25] [26] [27]. Since metabolomics analysis is often part of larger biomedical projects, there is an urgent need to standardize terminologies that cover sample preparation, experimental protocols, steps of software processing and metadata. While Asari fills a key gap in the computational metabolomics ecosystem, the fundamental issue of interoperable data structures remains a challenge. To standardize the computational aspects of metabolomics analysis and empower future computational developments, a set of common, well-defined, and reusable data structures will be essential, regardless of the programming language. This paper, therefore, describes a collection of common data structures involved in metabolomics data processing and illustrates how they are utilized in a full-featured Python-centric pipeline.

### Design and implementation

Semi-automated data analysis pipelines are essential for the mainstream adoption of metabolomics and its continued growth in the biomedical sciences. With pipelines, researchers of diverse backgrounds can process their data quickly and meaningfully, allowing for higher throughput and more extensive experiments. Furthermore, pipelines allow researchers to define highly reproducible workflows that are repeatable and reproducible by others. Our pipeline, named the Python-centric pipeline for metabolomics (pcpfm), enables start-to-finish metabolomics data processing based on Asari. The pipeline ingests centroided mzML data or Thermo raw files and returns a human-readable set of tables summarizing the detected features and their annotations and sample metadata. Annotation is a major step after preprocessing, utilizing multiple sources, such as authentic compound libraries and tandem mass spectral libraries. Annotation levels in pcpfm are described in accordance with Schmanski 2014 [28]. Additionally, the pipeline performs various processing steps, including normalization, feature interpolation, removal of rare features, quality assurance, quality control evaluations, and generates PDF reports to summarize results.

We designed a set of core data models, which are described in the MetDataModel package and summarized in Table 1. The goal of MetDataModel is to encourage reuse and extension, therefore the data models are kept minimal. Developers are free to extend them to more detailed and specific models. Such extensions and applications are exemplified here in the pipeline package, pcpfm.

**Table 1.** Core concepts implemented in the MetDataModel package.

| Name | Operational Definition |
| --- | --- |
| MS Spectrum | List of $m/z$ values and associated intensity, typically from a scan on a mass spectrometer |
| Mass Track | An extracted ion chromatogram of consensus $m/z$, spanning the full retention time. |
| Elution Peak | Peak of intensity values along the axis of chromatography. |
| Feature | A set of peaks that are aligned across samples, specific to an experiment. |
| Empirical Compound | A group of associated features, typically isotopes and adducts, that belong to the same tentative compound and co-elute if there is chromatography. |
| Compound | A metabolite or a chemical of xenobiotic origin, including contaminants. |
| Reaction | Biochemical process that interconverts one or more compounds, often catalyzed by an enzyme. |
| Enzyme | A protein that catalyzes a biochemical reaction. |
| Gene | An inheritable sequence of nucleotides, some of which code for proteins. |
| Metabolic Pathway | A series of linked reactions that typically involve structurally related compounds, usually defined by human knowledge. |
| Metabolic Network | A set of reactions connected by shared compounds. Mathematically identical to pathway, but not limited by pathway definition. |
| Metabolic Model | A collection of metabolic reactions and their associated metabolites, enzymes, and genes. Additional parameters, e.g. reaction rates and flux rates, can be included. |
| Study | A collection of experiments on a set of related samples. |
| Experiment | A set of acquisitions collected on a set of samples using consistent methods. |
| Method | The approach and parameters used for data collection in an experiment, e.g., chromatography and ionization parameters. |
| Sample | A biological sample or a control sample that is analyzed in a study. A sample can be analyzed in multiple experiments, by a single or multiple methods. An instance of data file generated by analyzing a sample is referred to as an acquisition. Analytical replicates need to be modeled explicitly if used. |

A mass spectrum typically consists of a list of m/z (mass to charge ratio) values and corresponding intensities. It can be from a full scan (MS^1^) or tandem mass spectrometry (MS^2^ and beyond). The mass spectrum can be in profile mode or centroid mode. In profile mode, the term “mass peak” is still used by some applications to refer to a group of m/z values that belong to the same ion species. Data in profile mode can be converted to centroid mode (mass peak picking) by software from the manufacturers or from scientific community, and usually done by default in format conversion to the common mzML format. Centroided data is much reduced in size and there is little reason to use profile mode.

A mass spectrometer is often connected to chromatography (typically liquid phase or gas phase); therefore, such an experiment acquires many mass spectra at different chromatographic retention times. Thus, data processing requires the detection of signals across spectra, i.e., scans. Such signals are typically presented as extracted ion chromatogram (EIC or XIC). In the Asari software, this concept of EIC is extended to a “mass track” [1], which is a vector of intensity values spanning the full scan range under one consensus m/z value. The use of mass tracks leads to new algorithms of alignment and feature detection [1]. Because “peak picking” or “peak detection” could refer to either mass peaks or elution peaks, we recommend the explicit term of elution peak detection. An elution peak is defined by ion intensity along the axis of retention time in the 2-dimensional representation. A mass peak is defined by ion intensity along the axis of m/z, usually in profile data. We define an elution peak at the level of a sample and as a feature at the level of an experiment (Table 1). The definition of “feature” here is consistent with its use in XCMS [12] and MZmine [29], but different from OpenMS [30]. OpenMS refers to a feature as a group of ions, likely due to its root in proteomics. The relationships between these concepts are illustrated in Figure 1A.

**Figure 1:**
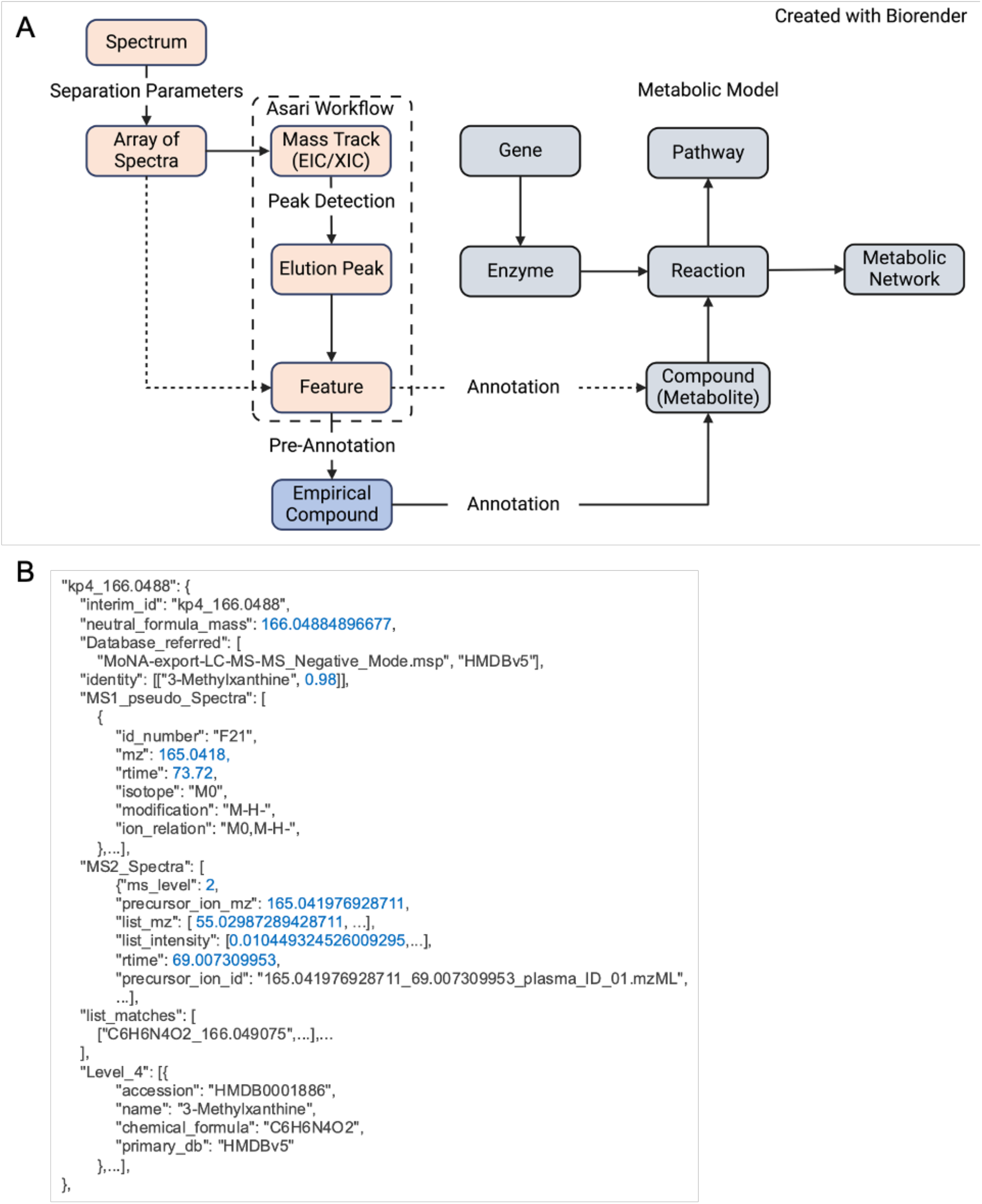
Design of core concepts and data models in computational metabolomics. **A)** The core concepts in MetDataModel, metabolomics data processing in salmon and metabolic modeling in grey. We introduce “empirical compound” as a key bridge in between. The dashed lines indicate alternative workflows. **B)** Abridged empirical compound example listing of MS^1^ features, annotation from MS^2^, and other sources. This JSON format enables chaining of multiple annotation tools.

The relationship between metabolite, reaction, enzyme, gene, pathway, and network is described on right side of Figure 1A, which are collectively considered as a “metabolic model”. Metabolic reactions are central to connect these entities, and the links to enzymes (proteins) and genes (measured in transcriptomics, genomics and epigenomics) are the most important basis for analyzing multi-omics data [31] [32]. These concepts mirror the extensive development from the field of genome-scale metabolic models (GSMMs) in over two decades. Connecting GSMMs with the experimental measurement by mass spectrometry is not trivial, because a) the identifiers of metabolites need to be consistent; b) charge states of molecules and experimental measurements need to be consistent; c) a significant knowledge gap exists between the GSMMs and experimental metabolomics; and d) metabolite identification is limited in experimental metabolomics.

The reality of metabolomics is that many features are not definitively identified. We have introduced the concept of empirical compound to describe the measurement of a tentative metabolite (Figure 1B). For example, in LC-MS (liquid chromatography coupled mass spectrometry) metabolomics, some isomers (molecules of identical mass) may not be resolved, limiting the annotation level. That is, the isotopologues and adducts clearly belong to the group, but the group may be isomer A, isomer B, or a mixture of both. Empirical compounds model this property and serve as an operational unit to link computational steps. It has been part of the software implementation since version 2 of mummichog and version 4 of MetaboAnalyst [33]. This design enables an organized presentation of degenerate MS^1^ features, and chaining annotation from MS^n^ and multiple methods. The isotopes and adducts from pre-annotation are modeled as a grid structure, made computable by the khipu package [34], which is also incorporated in the pcpfm pipeline. Annotation remains the most critical step in the meaningful interpretation of metabolomics data and the field faces the challenge of handling annotation uncertainty and probability. Empirical compounds provide an operational data structure as a path forward.

The abstract concepts in Table 1 and Figure 1 are intrinsically agnostic to programming languages. We demonstrate their implementation in Python 3 and JSON in MetDataModel. The pcpfm package is written in Python 3 and JSON is used extensively for intermediary data. Many pipeline data structures inherit from, and expand upon, objects provided by the MetDataModel library. Specific extension of empirical compound is exemplified in Figure 1B.

The inputs to our pipeline minimally consist of .mzML or .raw files and a metadata CSV file, that minimally maps sample names to acquisition file paths. While Asari was initially developed for orbitrap data, pcpfm is expected to be compatible with the data from major manufacturers that can be converted into mzML format [35]. The final output consists of a feature table detailing the observed m/z and retention time values for observed features mapped to unique identifiers, an annotation table mapping these identifiers to annotations and metadata for those annotations, and a third table summarizing the acquisition and experiment-level metadata. This three-table format handles multiple annotations gracefully and will be supported in future versions of MetaboAnalyst and Mummichog for downstream analysis and interpretation.

Each step in an analysis corresponds to one command in the CLI and one function in the main pipeline process (Supplemental Table 1). In brief, every analysis starts with assembling an experiment object from the metadata and acquisition data. This experiment object records the location of intermediates on disk for reuse in later steps. Optionally, any .raw files are converted to centroided .mzML files using the ThermoRawFileParser [36] before preprocessing with Asari which yields a “preferred” and “full” feature table.

Quality control is necessary in every project but depends on the experimental design. Multiple QA/QC operations are available including PCA, t-SNE, correlation cluster maps, various statistical tests that quantify sample properties such as feature count and median correlation, blank masking, the removal of outliers and uncommon features, normalization using median sample TICs (total ion counts), and missing value imputation. Some operations can be made batch aware and explicit batch correction is provided using pycombat [37] [38]. These operations are implemented using a mixture of Sklearn [39], Scipy [40] and Numpy [41], while Pandas [42] is used for generic data wrangling.

Empirical compounds can be constructed from a feature table using khipu [43] and most methods for empirical compounds concern annotation. Using MatchMS [44] [45], MS^2^ based annotations can be generated using data from DDA or deep scan workflows such as AcquireX [46] and MS^2^ spectral databases such as MoNA [47] or authentic standards libraries. MS^1^-based annotations are generated using our JSON metabolite services library and appropriately formatted inputs or *m/z* and retention time similarity to authentic standards. These annotations can be mapped back to any feature table to generate the previously mentioned tabular output. PDF reports can be created using the fpdf library [48] that summarize various intermediate results and records a timeline of the commands used for the analysis. Example reports are provided in Supplemental Files S2 and S3.

Most operations in the pipeline are chainable meaning they can be performed in a dynamic order with outputs from previous iterations being used as inputs. This flexibility allows users to build their workflows; however, example workflows are provided as .sh and nextflow scripts [49]. Nearly all parameters are user-configurable, but reasonable defaults are provided and documented, allowing the pipeline to be as hands-off or hands-on as the end user desires.

## Results

The pcpfm is designed to prepare data for downstream data analysis, which can be performed by bioinformaticians or data scientists without a background in mass spectrometry. The major steps are shown in Figure 2A. We demonstrate first the results on data processing, annotation, and quality control, then on biological applications. Seven metabolomics and one lipidomics datasets from four studies, three fully public and one in-house, are used in these examples (details in Supplementary File S1).

**Figure 2:**
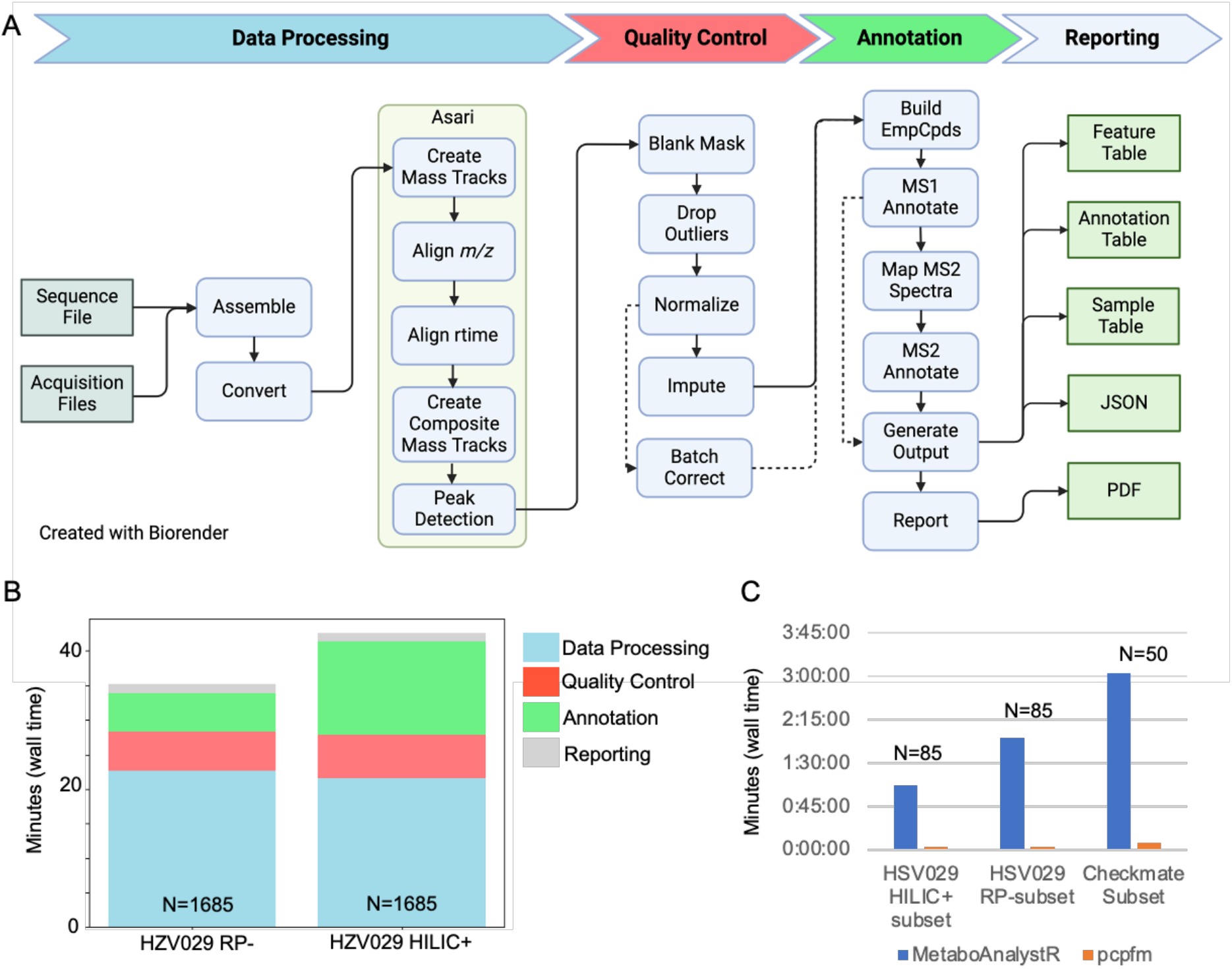
Design and computational performance of the pcpfm pipeline. A) The pipeline has four major sections: data processing, quality control, annotation and reporting. Data processing encapsulates everything from the start of a processing job to the creation of a feature table using Asari. Quality control consists of multiple chainable commands that allows for a raw feature table to be curated into a table suitable for downstream analysis. Annotation concerns the mapping of empirical compounds to metabolites using formula or MS2 similarity to databases, m/z and retention time mapping to authentic standards and optionally, MS2 similarity. Finally, reporting handles the creation of the three-table format for downstream analysis, PDF report generation, and JSON outputs for advanced users. Dashed lines represent common alternative workflows. B) Using the two largest datasets (n=1685), the high computational performance of our pipeline is demonstrated. Most of the wall time is spent during Asari. All steps are single threaded by default except Asari which uses 4 processes. Report generation excluded. C) A comparison of the wall time required for a minimal pcpfm workflow compared to its MetaboAnalystR equivalent.

A distinct advantage of pcpfm and Asari is the computational efficiency to process large datasets. The computational times are summarized on two high-resolution LC-MS datasets of 1685 samples. Processing and QC use less than half an hour on a laptop computer for each datasets, while the annotation step depends on the databases involved (Figure 2B). The computational performance of pcpfm is further compared to an XCMS-based workflow on a subset of three datasets (Figure 2C), showing clear improvement in our pipeline.

The metabolomics community have a consensus that metabolite annotation should be reported according to its confidence level. We have incorporated empirical compounds into both MS^1^ and MS^2^ annotations. By building empirical compounds first, i.e. pre-annotation via the khipu package, MS^1^ annotation is improved because the search of databases does not query many degenerate features (Figure 3A). The MS^2^ annotation utilizes MatchMS but with an optimization using an interval tree algorithm [50] . Because there are many implementations of MS^2^ annotation under similar principles, it is important to be explicit on the algorithm in pcpfm (Figure 3B). The MS^2^ annotation in pcpfm is efficient enough to run large experiments on consumer-grade hardware, as shown in Figure 2B. When authentic compounds are used to annotate metabolites, it is straight forward to match their m/z and retention time to biological samples (Figure 3C). Multiple annotations of different sources are chained in the empirical compound data structure (Figure 1B), which is amendable to future enhancements, e.g., context specific databases.

**Figure 3:**
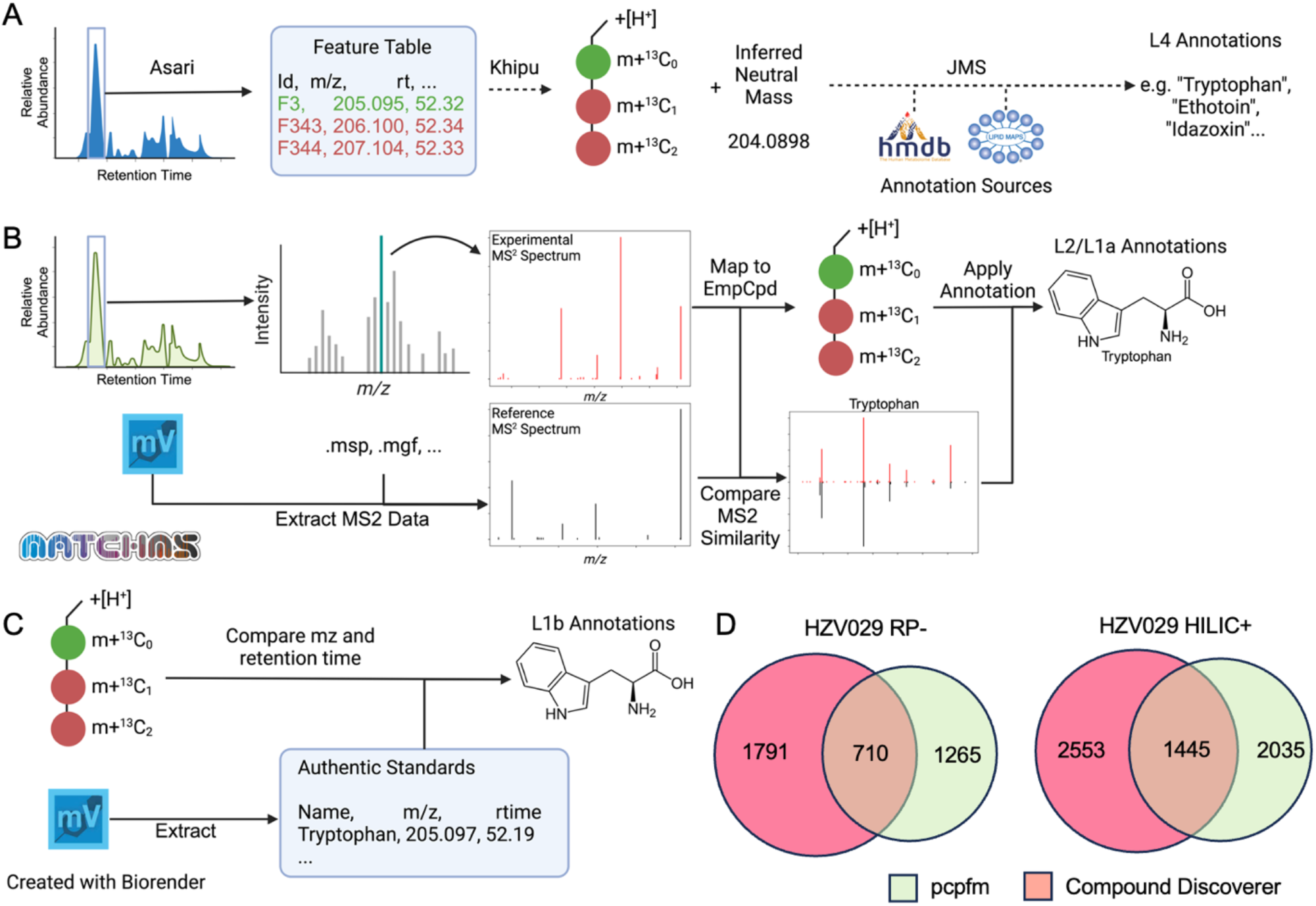
Annotation methods in pcpfm. **A)** Empirical compounds are constructed from Asari feature tables using khipu, which groups degenerate features such as isotopologues and adducts. The inferred neutral mass of an empirical compound is compared to known metabolites to generate level 4 annotation (via JMS, https://github.com/shuzhao-li-lab/JMS). **B)** Level 2 and 1a annotations are generated using MS^2^ similarity. Experimental MS^2^ spectra are mapped to empirical compounds and then compared to reference spectra, to annotate metabolite structures. **C)** Level 1b annotations are generated based on *m*/*z* and retention time match to authentic chemical standards. The use of empirical compound improves search efficiency and reduces false positives, while annotations at all levels can also be mapped to the feature level. **D)** Overlap of MS^2^ annotations by pcpfm and CD in the two HSV029 plasma datasets. Detailed dissection of the differences is difficult since CD is closed-source.

We compared the MS^2^ annotations generated by the pcpfm to those from vendor’s software, Compound Discoverer (CD). MS^2^ annotations were mapped back to the full Asari feature tables using an *m/z* tolerance of 10ppm and a retention time of 30 seconds, since the MS^1^ and MS^2^ experiments were performed separately. Considerable overlap is seen between CD and pcpfm annotations (Figure 3D). Because the algorithm in CD is closed source, it is not feasible to trace the differences between the tools, which highlights the importance of open-source tools for continued improvement.

The applications of pcpfm to quality control are demonstrated on a dataset consisting of 17 batches and 1685 samples (Figure 4). Multiple QC metrics can be plotted from Asari in the pipeline, so that users have a first-level visualization of data quality (Figure 4A). In this particular study, a recalibration of instrument occurred between batches 7 and 8, and the batch effect is revealed by inter-sample correlation (Figure 4B). Plots of TICs are useful for inspecting abnormal samples. With normalization and batch correction options in pcpfm, TIC plots show clear correction in the data (Figure 4C). The batch effect and correction is better illustrated by PCA plots (Figure 4D). Another common data quality issue is failed injections. Using Z-score metric of the number of features, pcpfm can detect both real and simulated failed injections (Figure 4E). These failed injections can be further verified by total ion chromatograms (Figure 4E).

**Figure 4:**
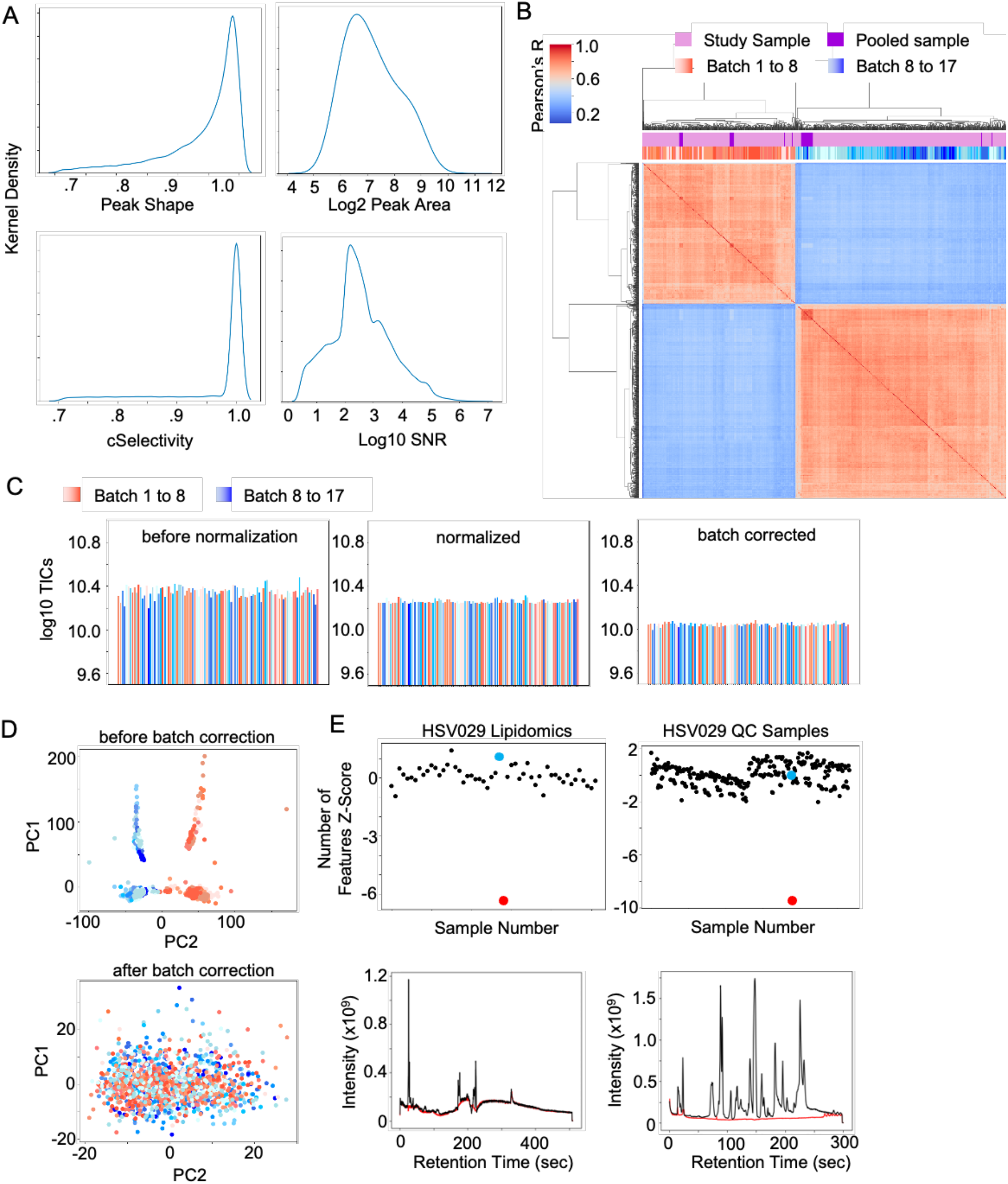
Examples of quality control in the pcpfm pipeline. **A)** A collection of QA/QC metrics generated by Asari on an example dataset (“HZV029 Plasma RP-”). **B)** The correlation clustermap of all study samples and pooled samples from the HZV029 Plasma RP-dataset (preferred feature table) illustrating the batch effect induced by instrument calibration. **C)** Log10 TICs of a random subset of samples before normalization, after normalization, and after batch correction. **D)** PCA demonstrating the presence of a batch effect (top) and its removal (bottom). **E)** Detection of failed acquisition by the number of feature Z-scores. The failed injection is highlighted in red and a representative “good” injection in blue for both the plasma lipidomics and HZV029 QC dataset (right and left, top). The lipidomics failed injection is simulated by replacing a missing sample with an empty vial while the other was identified post-hoc. The TICs of the failed and good injections are shown in red and black respectively (bottom).

Multiple previously published datasets were reanalyzed using pcpfm to evaluate the pipeline’s general suitability. Bowen et al (2023 [51]) designed a specialized xenobiotic-focused workflows to detect metabolites of the drug sunitinib. Our pipeline with default parameters detects all but one of the previously reported sunitinib-related metabolites in cardiomyocyte cell pellets and all features in culture media (Figure 5A). The sole missing feature is due to low signal-to-noise ratio, not passing Asari quality threshold (Figure 5B). The treatment of cardiomyocytes by sunitinib also induced a metabolic response [51], which is readily captured using ANOVA (Figure 5B). These results indicate the potential of pcpfm as a simplified yet broadly applicable workflow. To compare pcpfm feature detection against a state-of-art R-based pipeline (MetaboAnalystR), we reprocessed a subset of published metabolomics data on the CheckMate immunotherapy cohort [53]. The authors’ in-house metabolite library serves as a proxy of ground truth here. The result indicates that pcpfm detects more true metabolites (Figure 5C).

**Figure 5:**
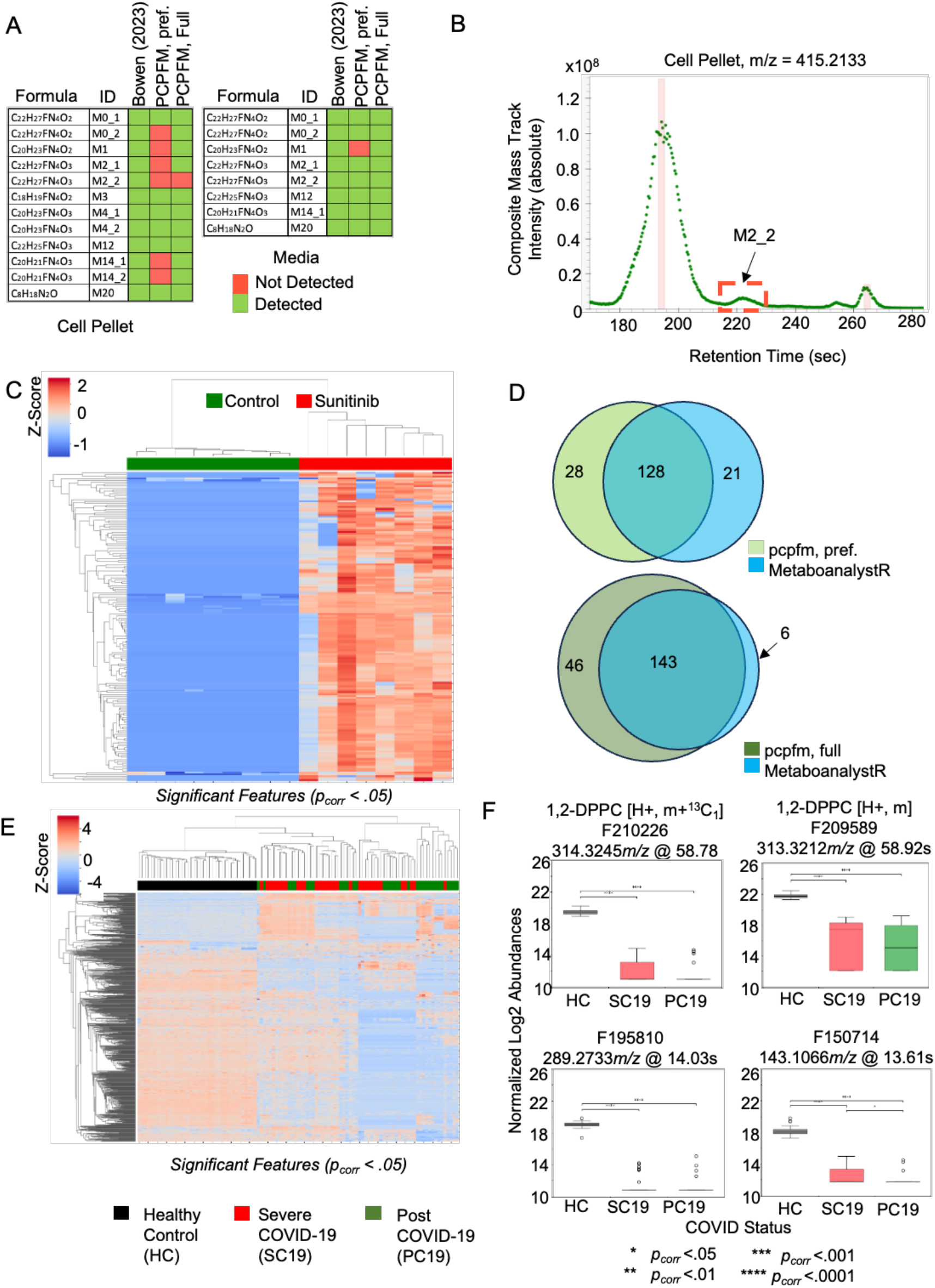
Applications of pcpfm to analyzing biological datasets. **A)** In Bowen 2023 dataset on cardiomyocytes, the pcpfm identifies most of the reported sunitinib-related features in both cell pellets and media using a standard workflow. Asari and pcpfm output both a preferred feature table and a full feature table, the former of higher feature quality and the latter more inclusive. **B)** The mass track for the sole feature undetected in the Bowen 2023 cell dataset is shown and the suspected undetected peak is in red box (M2_2), which fails to pass Asari quality requirement. **C)** Significant differential metabolite features between sunitinib exposure groups in cell pellets. ANOVA p-values are corrected for multiple testing by Benjamini-Hochberg method [52]. **D)** Both the pcpfm and MetaboAnalystR were used to extract features from a subset of the CheckMate study. Of 202 compounds in their authentic standard library, MetaboAnalystR identified 149, while the full table from the pcpfm identified 189 of the confirmed features. **E)** Clustering pattern of the Ansone 2021 cohort using features differentially abundant between treatment groups. **F)** Example boxplots of differentially abundant features in the Ansone 2021 cohort. F210226 and F209589 (top) were mapped to the same empirical compound that was tentatively annotated as 1,2-DPPC, a pulmonary surfactant by its sole level 4 annotation. Significance was evaluated using ANOVA and post-hoc Tukey’s HSD test in E and F.

Lastly, as an example for generating biologically meaningful results, we reanalyzed the metabolomics data from a COVID-19 exposure and recovery cohort (Ansone 2021, [54]). Following pcpfm, the significant features tested by ANOVA followed by a Tukey’s HSD [55] were subjected to hierarchical clustering (Figure 5E), which recapitulated the original observation that metabolic profile cluster by COVID infection and recovery vs. control in the Anosne 2021 paper. The box plots of selected features confirm the patterns of abundance changes in participant groups (Figure 5F). Interestingly, two features (Figure 5F, top) are found to belong to an empirical compound with a single level 4 annotation to 1,2-dipalmitoylphosphatidylcholine (1,2-DPPT), a pulmonary surfactant known to be less abundant in COVID patients than healthy controls [56]. This result demonstrates that novel biology can be gained with the pcpfm. The Jupyter notebooks underlying these examples are included in the pcpfm code repository, so that users can easily perform their own data analysis based on the templates.

### Availability and Future Directions

The MetDataModel and pcpfm are available through GitHub (https://github.com/shuzhao-li-lab/metDataModel and https://github.com/shuzhao-li-lab/PythonCentricPipelineForMetabolomics), and both are installable by pip via PyPi or from source. All dependencies are open source and downloadable via pip, except for the ThermoRawFileConverter and mono framework, both of which are optional. Example workflows are provided in bash and as nextflow; however, users can implement their own using the CLI or the pipeline internals available using standard Python conventions for APIs. API usage will be officially supported in an upcoming release.

Future development of pcpfm will implement additional options and methods for data processing, including normalization, interpolation, and batch correction. Improving support for non-orbitrap instruments is another priority for the pipeline and the underlying Asari algorithm. A cloud-based application is planned to allow users to process data in a friendly web interface.

## Data Availability

The version of the pcpfm, notebooks, and previously unreleased datasets used to generate the results presented in this manuscript are available at https://doi.org/10.5281/zenodo.10642162. The Checkmate dataset was retrieved from Metabolomics Workbench (https://www.metabolomicsworkbench.org) with study ID: ST001237. Bowen 2023 dataset was retrieved from Metabolights (https://www.ebi.ac.uk/metabolights/, accession code MTBLS2746). The Ansone 2021 dataset was retrieved from Metabolights (accession code MTBLS3852).

## Author Contributions

Conceptualization: JMM, SL; Data Curation: JMM, SL, YC; Formal Analysis: JMM, SL; Funding Acquisition: SL; Investigation: JMM, MT, ZP, JX, SL; Methodology: JMM, MT, JX, SL; Project Administration: SL, Resources: JX, SL; Validation: JMM, ZP, JX, SL; Visualization: JMM, SL; Writing – Original Draft Preparation: JMM, MT, MG, SL; Writing – Review and Editing: JMM, MT, YC, ZP, JX, SL.

## Acknowledgements

We would like to thank Paul Robson, Arti Taggar, Julianna Alcoforado Diniz, Zukai Liu, and Lucas Chang who graciously provided data for the initial development and testing of the pipeline. This work was in part supported by NIH grants U01 CA235493 (NCI), R01 AI149746 and AI149746 S1 (NIAID), and UM1 HG012651 (NHGRI).

## Supplemental Information

Supplemental Table 1. List of commands in the pcpfm pipeline, their inputs and outputs, and if they are chainable.

Supplemental File S1: Description of datasets, methods for generating previously unpublished datasets, and compound discoverer annotation workflow.

Supplemental File S2: HZV029 Plasma HILIC+ example PDF report.

Supplemental File S3: HZV029 Plasma RP-example PDF report.

Supplemental File S4: .zip of pcpfm v1.0.2

Supplemental File S5: .zip of MetDataModel v0.6.0

